# DyAb: sequence-based antibody design and property prediction in a low-data regime

**DOI:** 10.1101/2025.01.28.635353

**Authors:** Joshua Yao-Yu Lin, Jennifer L. Hofmann, Andrew Leaver-Fay, Wei-Ching Liang, Stefania Vasilaki, Edith Lee, Pedro O. Pinheiro, Natasa Tagasovska, James R. Kiefer, Yan Wu, Franziska Seeger, Richard Bonneau, Vladimir Gligorijevic, Andrew Watkins, Kyunghyun Cho, Nathan C. Frey

**Affiliations:** Prescient Design, Genentech, South San Francisco, CA, USA; Department of Antibody Engineering, Genentech, South San Francisco, CA, USA; Department of Structural Biology, Genentech, South San Francisco, CA, USA; Center for Data Science, New York University, New York, NY, USA; Department of Computer Science, Courant Institute of Mathematical Sciences, New York University, New York, NY, USA; Canadian Institute for Advanced Research, Toronto, Ontario, Canada

## Abstract

Protein therapeutic design and property prediction are frequently hampered by data scarcity. Here we propose a new model, DyAb, that addresses these issues by leveraging a pair-wise representation to predict differences in protein properties, rather than absolute values. DyAb is built on top of a pre-trained protein language model and achieves a Spearman rank correlation of up to 0.85 on binding affinity prediction across molecules targeting three different antigens (EGFR, IL-6, and an internal target), given as few as 100 training data. We employ DyAb in two design contexts: as a ranking model to score combinations of known mutations, and combined with a genetic algorithm to generate new sequences. Our method consistently generates novel antibody candidates with high binding rates, including designs that improve on the binding affinity of the lead molecule by more than ten-fold. DyAb represents a powerful tool for engineering therapeutic protein properties in low data regimes common in early-stage drug development.

## Introduction

Labeled data scarcity is foremost among the challenges for applying machine learning to biologic drug development. While among the best selling pharmaceuticals worldwide^1,2^, data generation for biologics is particularly costly and laborious. Such datasets often contain inherent noise and variation across batches^3^ and require careful quality control. Learning from sparse, noisy datasets is thus an ongoing challenge in early-stage drug discovery and lead optimization^4^. Moreover, the ‘best’ assessed candidate at early stages may need to be discarded down the line if a significant liability is discovered – e.g., chemical or physical instabilities, poor pharmacokinetics, or immunogenicity^4–6^. An overarching objective within this paradigm is thus to produce and refine a select cohort of the most promising therapeutic candidates as quickly and cheaply as possible.

Distinguishing between *differences* in measurements, rather than merely focusing on absolute values, offers potential to address data scarcity. For example, Fralish et al. developed DeepDelta^7^, a deep learning model that predicts differences in pharmacokinetic properties between pairs of small molecules. By learning on concatenated representations of molecular structure pairs, their method showed good performance on small datasets containing ~ 200-1000 labeled points. A pair-wise method can provide multiple benefits, including preventing over-fitting and batch effects by combinatorially expanding the training dataset and potentially improving out-of-distribution performance. Rather than learning on *N* raw values, a model would instead learn on 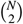 pairs. For example, a dataset of 500 points would swell to one containing nearly 125,000 pairs of data. Here, we show that a similar approach can be applied to large molecules by leveraging embeddings from language models trained on protein sequence data. Such protein language models (pLMs) have shown recent success on property prediction and engineering tasks, learning powerful representations of protein sequences^8–13^.

Our methodology captures the nuances of protein sequence variation by learning on relative embeddings and property differences, rather than absolute values. We call our model ‘DyAb’ - from **Dy**ad, meaning a pair, and **A**nti**b**ody, the protein class on which we focus here. One of the most important steps in antibody development is *in vitro* affinity maturation^14^. This process can entail screening single point mutants for binding against the antigen of interest, and selecting variants with improved affinity. We employ our model on three such datasets from three different targets (i.e., disease programs), containing as few as 100 labeled points. For each set, we generate novel antibody sequences that express in mammalian cells and bind strongly to the target of interest. Moreover, our method can design mutation combinations and antibody variants with improved affinity on the single point mutants. DyAb thus represents a promising approach for sequence diversification and optimization in the early stages of biologic therapeutic development, where labeled data can be few and far between.

## Results

### Learning differences between protein pairs in a low-data regime

The DyAb model framework was inspired by the types of datasets common in early-stage biologic development – specifically, data from COmprehensive Substitution for Multidimensional Optimization (COSMO) experiments^15^. Such experiments mutationally scan residues in antibody complementary-determining regions (CDRs) with all natural amino acids, except cysteine. The resulting dataset contains affinity labels for ~ 500-1000 point variants around a lead molecule, providing insights into the most important residues for antigen binding. Our model leverages these closely-related variant sets to learn relationships between antibody pairs and predict novel, higher edit distance sequences that maintain antigen binding. For example, in this pair-wise framework, the model learns on almost 125,000 pairs of binding data rather than the 500 point mutant data in the COSMO set.

DyAb takes pairs of protein sequences as input (Fig. 1a). Protein sequences are first fed through a pre-trained protein language model (pLM), from which embeddings are extracted (Fig. 1b). Here, we primarily discuss results using AntiBERTy^16^ and LBSTER^17^, pLMs trained exclusively or primarily on antibody sequences. We then compute a ‘relative embedding’ for each unique sequence pair by taking the difference between their embeddings. Please see Supplementary Fig. S1-S2 for more detailed descriptions of this framework. A convolutional neural net (CNN) trained on these relative embeddings and property labels then outputs a prediction of the property difference between two sequences (Fig. 1c). Thus, DyAb can either be used as a ranking model to score combinations of known mutations, or paired with a sampling algorithm to directly generate new sequences (Fig. 1d). Both are explored in this work.

**Figure 1.**
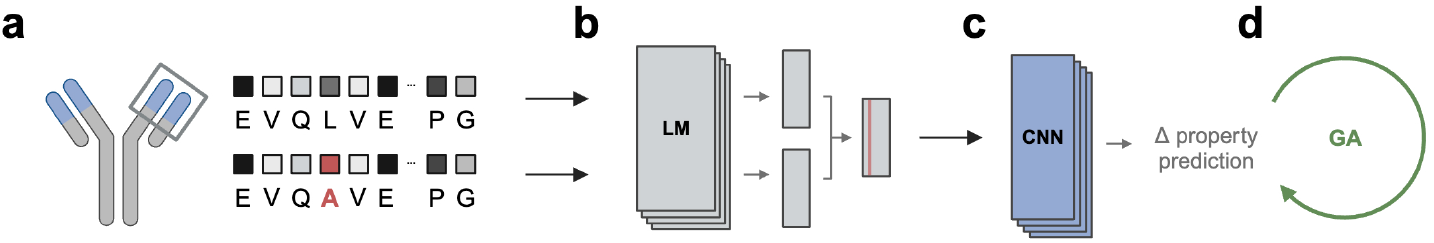
Overview of the DyAb workflow for protein property prediction. **(a)** Pairs of closely-related protein sequences – here, the variable domain of a therapeutic antibody (blue) – are fed through a **(b)** pre-trained language model (LM). The relative embedding between these sequences is then **(c)** used as input to a convolutional neural network (CNN) to predict differences in properties of interest, namely binding affinity. **(d)** A genetic algorithm (GA) is optionally employed to sample novel mutation combinations.

We tested DyAb on three antibody-antigen binding affinity datasets of varying size and complexity (Fig. 2). Here, binding affinity is defined as the log transform of the equilibrium dissociation constant, p*K*_D_ = *−* log_10_(*K*_D_) = −log_10_(*k*_d_*/k*_a_), where *k*_d_ and *k*_a_ are the measured dissociation and association rate constants, respectively. Each set contains labels for variants around an antibody ‘lead,’ a known binder of a given antigen. Binding affinity differences, Δp*K*_D_, are then computed relative to this lead. The first dataset represents the use case that inspired the model: a COSMO dataset of *N* = 512 point mutants of an internal antibody (anonymized here, termed ‘lead A’). DyAb performs well on a held-out test set of *N* = 77 point variants (*N* = 2926 pairs), with a Pearson correlation coefficient *r* = 0.77 and Spearman rank correlation coefficient *ρ* = 0.80 (Fig. 2a, p < 0.001 for both).

**Figure 2.**
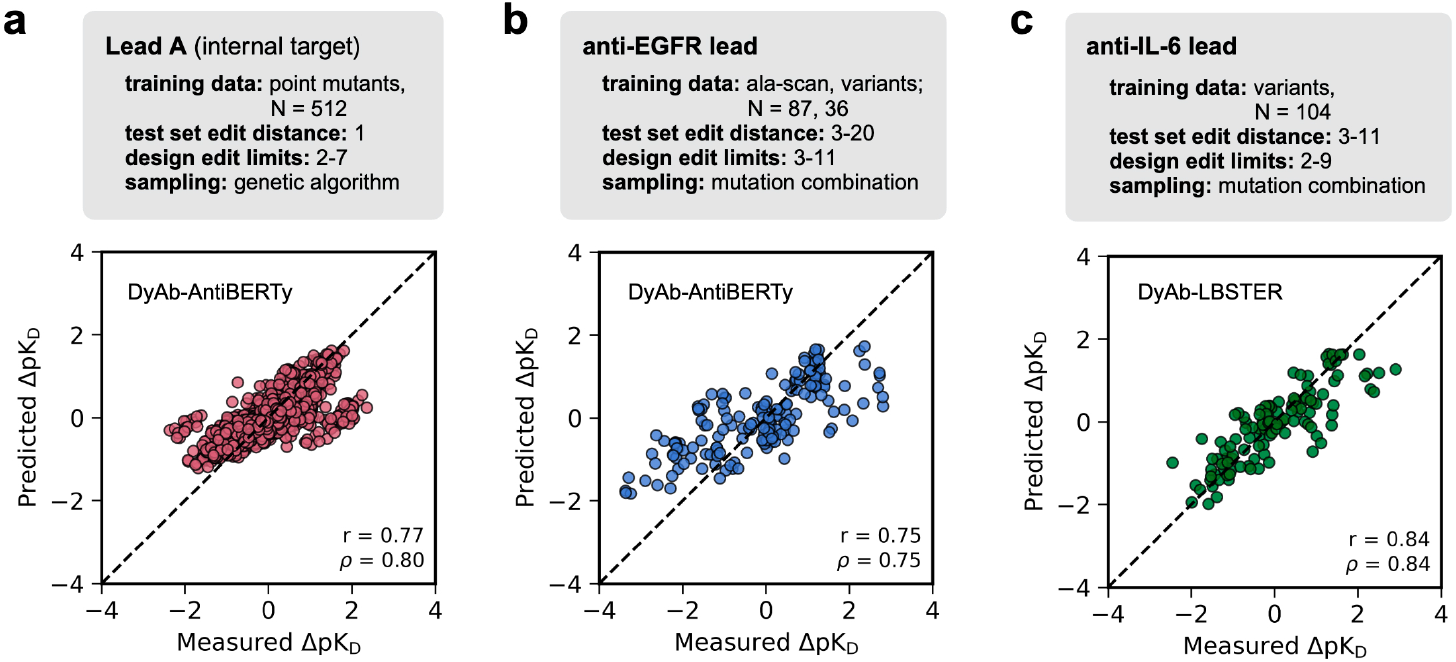
DyAb model performance on small antibody affinity datasets. Predicted versus measured improvements in affinity, ΔpK_D_, for variants of three lead antibodies (upper, colors, **(a-c)**), for the test set held-out during training. Pearson (*r*) and Spearman (*ρ*) correlation coefficients are reported for each test set.

We also evaluated DyAb on more difficult data landscapes for anti-EGFR and anti-IL-6 leads, given only ~100 affinity labels for each. These sets included more distant variants of the lead, within 20 edit distances (ED, see Supplementary Fig. S4). Such variants might typically be obtained via rounds of *in silico* or *in vitro* affinity maturation^14^. When trained only on point-variant data from alanine-scanning mutagenesis^18^ and a handful of higher ED variants, DyAb predicts variant Δp*K*_D_ with *r* = 0.75 and *ρ* = 0.75 (Fig. 2b; p < 0.001 for both). We observed good performance even in the complete absence of point-variant data, with *r* = 0.84 and *ρ* = 0.84 over the anti-IL-6 variant test set (Fig. 2c; p < 0.001 for both).

### DyAb generates novel, expressing antibody variants with high binding rates

We next applied DyAb to generate novel variants of these three lead antibodies and further optimize antigen binding affinity. First, for the lead A set, we selected all COSMO mutations that individually improved affinity. We then generated all combinations of these mutations with ED = 3-4, and used the trained DyAb-AntiBERTy model (Fig. 2a) to score designs by predicted ΔpK_D_ relative to the lead. Starting from this initial population, we employed a genetic algorithm (GA) to select and mutate these sequences to sample the vast design space and iteratively improve the predicted ΔpK_D_ (see *Methods*). The 47 top-ranked designs produced by this DyAb-GA model were selected for experimental testing (Fig. 3a, termed R1). 85% of this design set successfully expressed in mammalian cells and bound to the target antigen, an improved binding rate to that of the COSMO point mutants (59%). DyAb performance on the regression task for design sets are shown in Supplementary Fig. S3. Of all DyAb-designed binders against target A, 84% improved on the parent affinity of 76 nM, with the strongest binder reaching 15 nM.

**Figure 3.**
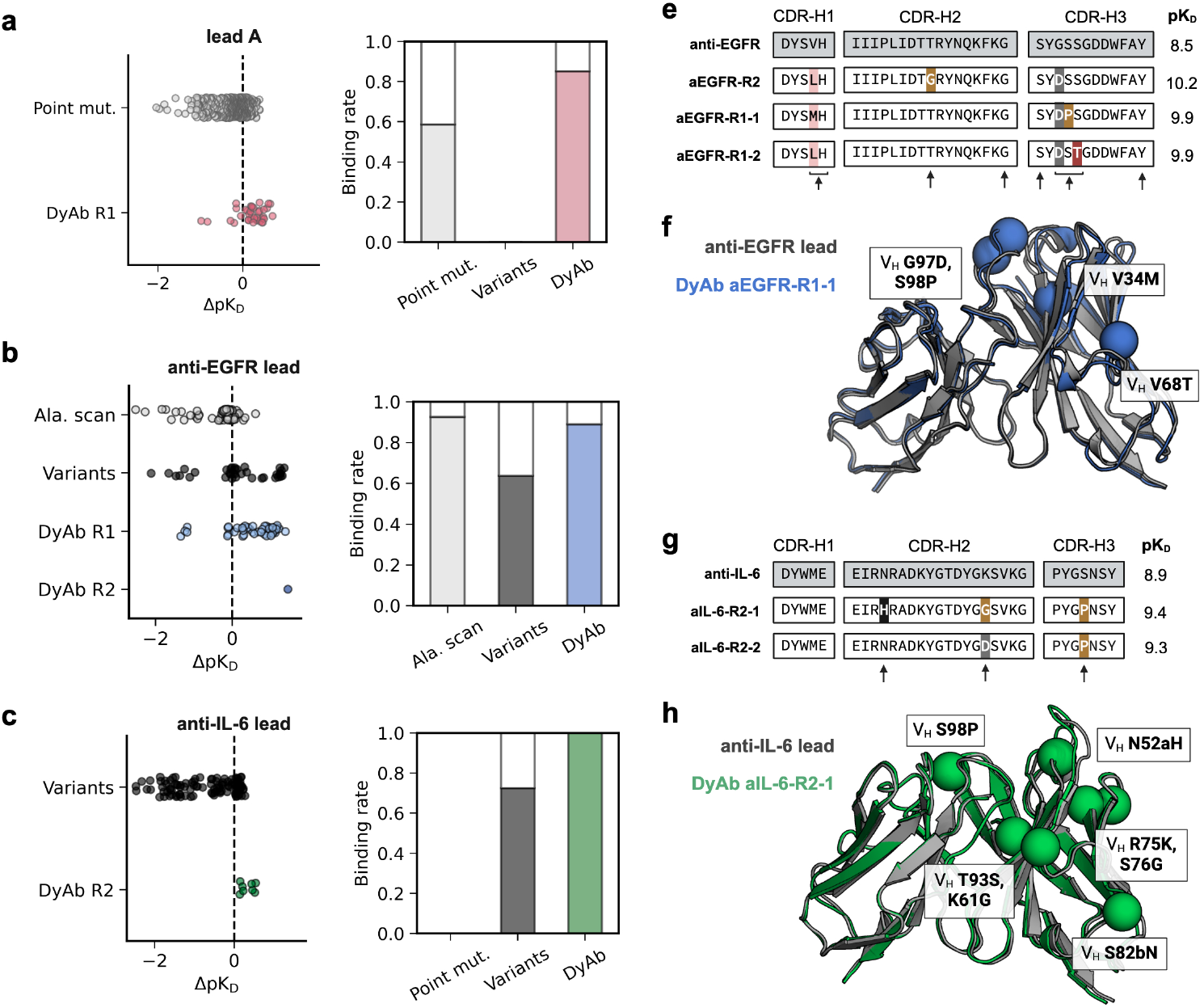
DyAb designs expressing antibody candidates against multiple antigens with high binding rates. **(a-c) (left)** Affinity improvements relative to the starting lead molecule, ΔpK_D_, for the training datasets of point mutations (light grey, general ‘point mut.’ or alanine-only ‘ala. scan’) and higher edit-distance variants (black), and DyAb-generated designs (colors). R1 models used AntiBERTy embeddings^16^ and R2 used LBSTER embeddings^17^. **(right)** Binding rates for antibodies in the training set and DyAb designs. **(e, g)** Sequence analysis of the heavy chain CDRs for the highest-affinity DyAb designs against EGFR and IL-6. Mutations are colored by amino acid character: aliphatic (pink), polar (red), negative (grey), positive (black), and other (gold). Arrows denote sites of all H-CDR mutations found in DyAb-designed binders. Yield is shown in units of mg/ml. **(f, h)** Structural analysis of DyAb designs and their starting leads. Anti-EGFR structures were solved experimentally (PDB entries *9MU1* for the lead and *9MSW* for aEGFR-R1-1), whereas Fv structures for the anti-IL-6 lead and R2-1 design were predicted via ABodyBuilder2^20^. Mutations in the CDRs and framework regions (spheres) are labeled.

We followed a similar procedure for designing anti-EGFR variants. First, we selected all mutations in the training set that improved affinity individually (in the alanine-scan) or within a higher ED variant. Instead of employing a GA, we then exhaustively generated and scored all combinations of these mutations between edit distances of 3 and 11. Designs with a DyAb-predicted ΔpK_D_ *>* 0 were then submitted to another ranking oracle^17,19^that selected a final 44 sequences for experimental testing. In the first round, 89% of designs expressed and bound EGFR (Fig. 3b, R1). 79% of these binders had stronger measured affinities than that of the starting point lead candidate (3.0 nM), including eleven designs with at least a ten-fold improvement. While the strongest binders had similar affinities to high-ED variants in the training set ( ~100 pM), DyAb produced binders at a much higher rate, on par with that of Alanine point mutants. We then undertook a second round of design that incorporated these data back into the training set. A single sequence was selected for experimental testing (R2, Fig. 3b). This design expressed, bound EGFR, and further improved affinity to 66 pM, exhibiting a near 50-fold improvement.

Finally, we tested if DyAb can generate antibodies with favorable properties given data for only ~ 100 variants of an anti-IL-6 lead. As before, we selected and exhaustively combined all mutations from affinity-improving variants in the training set up an ED of 9, and scored the generated sequences with a DyAb-LBSTER model (Fig. 2c). A subset of 8 designs with predicted improved affinity ΔpK_D_ *>* 0 were then selected^19^ for experimental testing (Fig. 3c, R2). All generated sequences were successfully expressed, bound IL-6, and improved affinity relative to the lead (1.4 nM). Four designs increased affinity by more than 3-fold, surpassing all variants in the training set.

### Structural analysis of top DyAb designs

While DyAb is a sequence-based model, analysis of top designs suggests putative structural mechanisms underlying the observed affinity improvement. We determined crystal structures of the corresponding Fabs of the anti-EGFR lead and R1-1 design to 2.4Å and 2.1Å resolution, respectively (Supplementary Fig. S7). DyAb makes four mutations in aEGFR-R1-1 across CDR-H1, CDR-H3, and the framework. Mutating Kabat position V_H_ 97 from Glycine to Aspartic acid – consistent across all top designs (Fig. 3e) – alters the conformation of CDR-H3. An additional Proline at V_H_ 98 could further stabilize this binding-able conformation in aEGFR-R1-1 relative to the lead. All three top designs also mutate V_H_ 34 at the base of CDR-H1 from Valine to a longer aliphatic amino acid.

In lieu of experimental structures, we compared variable domains predicted by ABodyBuilder2^20^ for the highest-affinity anti-IL-6 design and its lead (Fig. 3h). While the same V_H_ S98P mutation is seen in the top two designs, the predicted CDR-H3 conformation remains unchanged. Instead, a mutation in CDR-H2 (N52aH) drives extension of the loop into solution (Fig. 3g).

### Antibody-specific protein language models improve DyAb performance

DyAb’s pair-wise framework can be constructed from a variety of language model embeddings. While this work focused on antibody-specific pLMs, many general protein language models exist. Here, we compare DyAb performance head-to-head across all three variant datasets using embeddings from three different pLMs: AntiBERTy^16^ (trained on 500M natural antibody sequences), LBSTER^17^ (trained on the same antibody sequences, plus the general protein universe), and ESM-2^21^ (trained on just the general protein universe).

We evaluated DyAb models trained using relative embeddings from each pLM on the same train/test sets shown in Fig. 2. Performance on the downstream Δp*K*_D_ prediction task varies between datasets (Fig. 4). In some cases, we observed better performance with a different pLM than was used for design. For example, anti-IL-6 R2 designs were generated using LBSTER embeddings, which are slightly outperformed by AntiBERTy-based models across two of the four evaluation metrics. However, pLMs trained primarily or entirely on antibody repertoires produced more predictive DyAb models across the board versus a general pLM (ESM-2).

**Figure 4.**
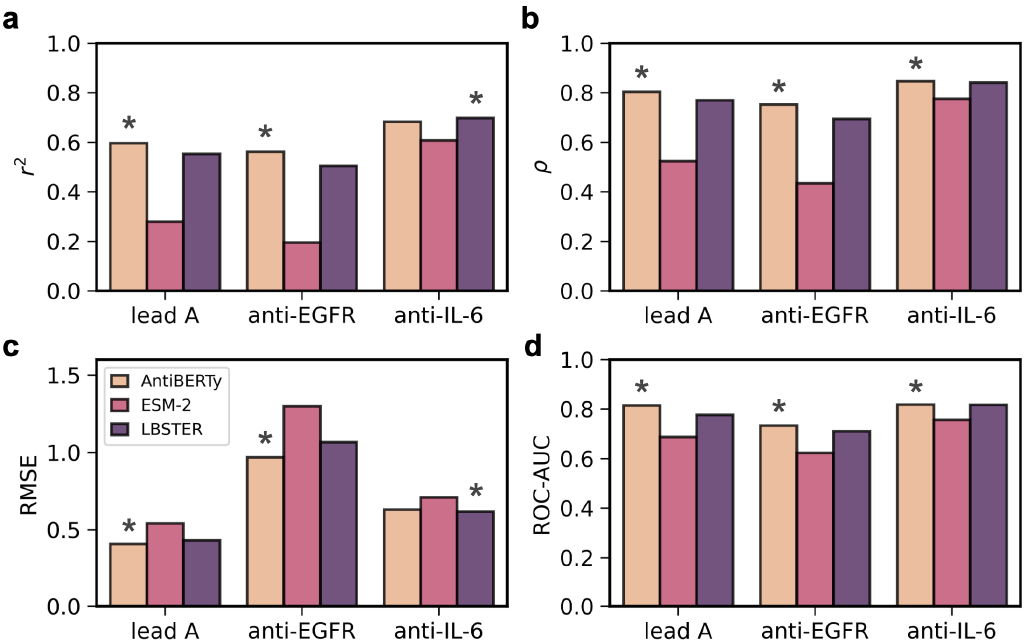
Ablation of language model embeddings. Evaluation of DyAb model performance on the held-out test set for all three antibody leads, trained with relative embedding from three pLMs: AntiBERTy^16^, ESM-2^21^, and a LBSTER pLM trained on antibody-specific and general data^17^ (see *Methods*). Asterisks denote the best-performing model on that dataset for each metric: **(a)** Pearson *r*^2^, **(b)** Spearman *ρ*, **(c)** root mean squared error RMSE, and **(d)** the area-under-the-curve (AUC) of the Receiver Operating Characteristic (ROC) curve when predicting a binary improved/worsened affinity label with a threshold of ΔpK_D_ = 0.

## Discussion

We developed DyAb, a deep learning model that leverages sequence pairs to predict protein property differences in a limited data regime. When applied to antibody design, DyAb efficiently generates novel sequences with enhanced properties given as few as ~100 labeled training data. Designs express and bind at consistently high rates (*>* 85%), comparable to that of single point mutants. Most DyAb-generated sequences improve upon the affinity of the lead and even upon that of the training set. DyAb thus represents a promising tool for early-stage antibody lead optimization and diversification.

We tested DyAb both as a ranking model for scoring combinations of mutations, and paired with a genetic algorithm (GA) for sampling. The main failure mode of GAs is a deviation from ‘natural’ sequences after long optimization trajectories away from the starting sequence, exceeding *~*8 edits^19^. We avoid this issue by setting a low edit distance design limit (ED = 7), incorporating only mutations found in previously stable sequences, and using pLM likelihoods in the discriminator. DyAb can be readily integrated with other algorithms like Monte Carlo tree search^22^ or generative methods like PropEn^23^ to further sample the design space. Future iterations could also incorporate protein structural features by leveraging embeddings from structure-informed models like ESMFold^24^ or SaProt^25^. Moreover, experimental structures for DyAb-designed antibodies in co-complex with antigens could provide additional insight into binding mechanisms and model rankings.

We note that the anti-EGFR and anti-IL-6 designs tested in experiment were sub-selected from a larger pool of top designs by another ranking oracle^19^, responsible for internal cross-project sequence prioritization. The designs against target A were selected solely based on DyAb scores. In all cases, DyAb was responsible for creating the enriched library containing the improved binders, regardless of what final ranking model was used.

The ability to learn in a low-N regime makes DyAb a promising method for engineering other antibody properties for which data are even scarcer. Chemical and physical stability at high concentrations are particularly important for drug development, but material-intensive to measure^6^. High-throughput proxy assays for these properties – e.g., that measure melting temperature^26^, self-interactions^27^, or solubility^28^ – could easily generate datasets of ~ 100 variants on which DyAb can learn well. While we focused on antibody therapeutics in this study, our model is general and can be applied to any protein variant dataset. DyAb is made publicly available as a model within LBSTER^17^.

## Methods

### Data preparation

For the regression task, we limited each affinity dataset to only antibodies that bind the target antigen. Here, a ‘binder’ is defined as having a measurable p*K*_D_ between 4 and 11. We filtered out noisy labels and those for which the SPR curves are poorly fit, as assessed by visual inspection. The datasets for lead A and anti-IL-6 were randomly divided into an 85%/15% train/test split. The anti-EGFR training set contained all alanine-scanning point mutants and a randomly-selected half of the higher-ED variants. The remaining half were held out for model testing.

### DyAb model framework

Antibody variable domain sequences were first aligned to the AHo-numbering scheme^29^ with ANARCI^30^ and concatenated to form variable domain sequences of consistent length (heavy followed by light chains). These sequences were then fed through and embeddings extracted from one of three pre-trained pLMs. AntiBERTy^16^ is a transformer-based masked language model, trained on over 500M natural antibody sequences from the Observed Antibody Space (OAS^31^). ESM-2^21^ is a general pLM trained on ~ 65M protein sequences from the UniRef50^32^ dataset. Here we employ the version trained with 8M parameters and 6 layers, termed *esm2_t6_8M_UR50D*. The LBSTER^17^ model employed here is a 11M parameter causal pLM trained on antibodies in OAS, general proteins from UniRef50, and protein complexes in the Protein Databank (PDB^33^) as of October 20, 2023. We chose this specific LBSTER model because it historically performed well on affinity-prediction tasks for internal Genentech antibodies.

Embeddings were then frozen, resized, normalized by the minimum tensor value, and rescaled based on the tensor maximum. Relative embeddings for each sequence pair {*s_i_*, *s_j_*} were obtained by element-wise subtraction of the normalized embeddings.

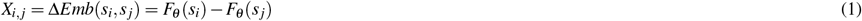

where *F*_*θ*_ is the protein language model. For the downstream task, these relative embeddings *X*_*i*, *j*_ were used as input for a CNN trained to predict the property difference between *s*_*i*_ and *s*_*j*_. We employed a ResNet18 architecture^34^ as implemented by PyTorch^35^ and an Adam optimizer^36^ with a learning rate of 2*e* − 4. Models were trained for 100 epochs using a mean-square error loss function and a batch size of 32. For the models shown in Fig. 2, we enumerated and trained on all unique pairs of molecules in the training set. For faster training, we also deployed a model that randomly sampled molecule pairs in the training set in each epoch. For inference, the relative embedding to the lead molecule *X*_*i*,lead_ was computed for a given pLM, *F*_*θ*_, and used to predict the property difference relative to the lead.

### Genetic algorithm

We employ a genetic algorithm (GA) for sampling when generating DyAb designs against target A. Inspired by the process of natural selection, GAs have found promising applications in the realm of protein design^37^. In this context, mutants can be viewed as variants of their parents. The fitness of each protein is evaluated based on the DyAb-predicted Δp*K*_D_ relative to the parent. Starting with an initial population of candidates, GA iteratively select, crossover, and mutate these sequences to explore the vast protein design space. Through successive generations, less favorable sequences are eliminated, and more optimal ones are retained and refined. We implement the following workflow:

- Start with a parental clone – here, lead A.
- Pick all mutations in the training set that individually improved binding affinity.
- Randomly select 3-4 mutations from this set and combine to generate new sequences.
- Score the new sequences with DyAb, *f* (newseq, oldseq), to get a predicted affinity difference Δp*K*_D_.
- Keep the most promising sequence as a new lead, and repeat the process.

### Protein expression and purification

Variable domains of DyAb designs were synthesized by IDT (Integrated DNA Technolgies), amplified using PrimeStar Max polymerase (Takeda), cloned into mammalian expression vectors using Gibson assembly, and transiently expressed in Expi293 cells by 1 mL cultures. After 7 days, cultured supernatants were harvested for antibodies purification as previously described^38^.

### Binding affinity measurements

Binding affinities of DyAb designs against their target antigens were determined by surface plasmon resonance (SPR) on a Biocore 8K machine (Cytiva). SPR was assessed at 37degC in HBS-EP+ buffer (10 mM Hepes, pH 7.4, 150 mM NaCl, 0.3mM EDTA and 0.05% vol/vol Surfactant P20). Designs against target A were assessed in a single cycle and those against EGFR and IL-6 in multi-cycle mode. Antibodies were captured on a Protein A chip, followed by injection of the target antigen for 5 minutes, dissocation for 10 minutes at 30 *µ*L/min, and regeneration of the surface with 10 mM glycine pH 1.5. The sensorgrams were then recorded and fit to a 1:1 Langmuir binding model to determine the equilibrium dissociation constant, *K*_D_. A log-transform produces the affinities reported in this work, p*K*_D_. Raw sensorgrams for the highest affinity DyAb design against each antigen are shown in Supplementary Fig. S5. Predicted affinity differences, Δp*K*_D_, are reported relative to the lead as measured in each experimental round. Absolute affinities for each lead (Fig. 3e,f) are reported as the median p*K*_D_ measured across all experimental rounds.

### Protein expression, purification, crystallization, and structure determination

Fab constructs for the aEGFR lead and R1-1 design were transfected into CHO cells at a 1:2 (HC:LC) DNA ratio, and expressed for 10 days. After harvesting, the supernatant was collected for purification using GammaBind Plus Sepharose (Cytiva) followed by size exclusion or SP cation exchange chromatography. The purified Fabs were formulated in 20mM histidine acetate pH 5.5, 150 mM NaCl.

Proteins were concentrated to 10-20 mg/mL and crystallized in [0.2M Ammonium sulfate, 0.1M sodium acetate pH 4.6, 30% w/v PEG 2000MME] for the aEGFR lead and [0.2M sodium bromide, 20% w/v PEG 3350] for DyAb design aEGFR-R1-1. Proteins were cryoprotected with the addition of 5-15% ethylene glycol.

Crystallographic data were collected at the Shanghai Synchrotron Radiation Facility (SSRF), beamline BL02U1. Data were integrated scaled with XDS^39^, and the structures were determined by molecular replacement with PHASER^40^. The model was built in COOT^41^ and subsequently refined with PHENIX^42^ to final statistics presented in Supplementary Table S1. The simulated annealing omit difference electron density maps (Supplementary Fig. S7) demonstrate the fidelity of the final models with the electron density.

## Figures

All figures were created with BioRender.com. Antibody structures were visualized with PyMol^43^.

## Acknowledgements

The authors thank the entirety of Prescient Design and the Genentech Antibody Engineering department for insightful discussions. We thank Yanshu Dou and Viva Biotech for structural work and the Shanghai Synchrotron Radiation Facility (SSRF) and staff of beamline BL02U1 for crystallographic data collection. We gratefully acknowledge the Prescient Design engineering team and Genentech IT for computing support.

## Author contributions statement

J.Y.L., A.L., K.C., and N.C.F. conceived the work; J.Y.L., J.L.H., A.L., S.V., E.L., P.O.P., N.T., R.B., V.G., F.S., A.W., K.C., and N.C.F. designed computational experiments. W.L. and Y.W. designed and conducted all wet-lab experiments, except the x-ray crystallography which J.R.K. designed and conducted. J.Y.L., J.L.H., A.L., W.L., S.V., E.L., P.O.P., N.T., F.S., R.B., V.G., A.W., K.C., and N.C.F. analyzed the results. J.L.H., J.Y.L., and N.C.F. wrote the initial draft and all authors reviewed the manuscript.

## Additional information

### Data availability

All data associated with this work are provided, excepting the sequences for the lead A dataset and designs, which are proprietary to Genentech. Coordinates and amplitudes for the crystal structures have been deposited at the RCSB with accession codes *9MU1* and *9MSW*. DyAb is implemented and available for use under an Apache-2.0 license within LBSTER^17^ at github.com/prescient-design/lobster.

## References

1. Lu, R.-M. et al. Development of therapeutic antibodies for the treatment of diseases. J. biomedical science 27, 1–30 (2020).

2. Kelley, B. Developing therapeutic monoclonal antibodies at pandemic pace. Nat. biotechnology 38, 540–545 (2020).

3. Wossnig, L., Furtmann, N., Buchanan, A., Kumar, S. & Greiff, V. Best practices for machine learning in antibody discovery and development. Drug Discov. Today 104025 (2024).

4. Akbar, R. et al. Progress and challenges for the machine learning-based design of fit-for-purpose monoclonal antibodies. In MAbs, vol. 14, 2008790 (Taylor & Francis, 2022).

5. Carter, P. J. & Rajpal, A. Designing antibodies as therapeutics. Cell 185, 2789–2805 (2022).

6. Zarzar, J. et al. High concentration formulation developability approaches and considerations. In MAbs, vol. 15, 2211185 (Taylor & Francis, 2023).

7. Fralish, Z., Chen, A., Skaluba, P. & Reker, D. Deepdelta: predicting admet improvements of molecular derivatives with deep learning. J. Cheminformatics 15, 101 (2023).

8. Ruffolo, J. A. & Madani, A. Designing proteins with language models. nature biotechnology 42, 200–202 (2024).

9. Dallago, C. et al. Flip: Benchmark tasks in fitness landscape inference for proteins. bioRxiv 2021–11 (2021).

10. Hie, B. L. et al. Efficient evolution of human antibodies from general protein language models. Nat. Biotechnol. 42, 275–283 (2024).

11. Li, L. et al. Machine learning optimization of candidate antibody yields highly diverse sub-nanomolar affinity antibody libraries. Nat. Commun. 14, 3454 (2023).

12. Hayes, T. et al. Simulating 500 million years of evolution with a language model. bioRxiv (2024).

13. Madani, A. et al. Large language models generate functional protein sequences across diverse families. Nat. Biotechnol. 41, 1099–1106 (2023).

14. Kennedy, P. J., Oliveira, C., Granja, P. L. & Sarmento, B. Monoclonal antibodies: technologies for early discovery and engineering. Critical reviews biotechnology 38, 394–408 (2018).

15. Sampei, Z. et al. Antibody engineering to generate sky59, a long-acting anti-c5 recycling antibody. PLoS One 13, e0209509 (2018).

16. Ruffolo, J. A., Gray, J. J. & Sulam, J. Deciphering antibody affinity maturation with language models and weakly supervised learning. arXiv preprint arXiv:2112.07782 (2021).

17. Frey, N. C. et al. Cramming protein language model training in 24 gpu hours. bioRxiv 2024–05 (2024).

18. Cunningham, B. C. & Wells, J. A. High-resolution epitope mapping of hgh-receptor interactions by alanine-scanning mutagenesis. Science 244, 1081–1085 (1989).

19. Gruver, N. et al. Protein design with guided discrete diffusion. Adv. neural information processing systems 36 (2024).

20. Abanades, B. et al. Immunebuilder: Deep-learning models for predicting the structures of immune proteins. Commun. Biol. 6, 575 (2023).

21. Rives, A. et al. Biological structure and function emerge from scaling unsupervised learning to 250 million protein sequences. PNAS (2019).

22. Świechowski, M., Godlewski, K., Sawicki, B. & Mańdziuk, J. Monte carlo tree search: A review of recent modifications and applications. Artif. Intell. Rev. 56, 2497–2562 (2023).

23. Tagasovska, N., Gligorijević, V., Cho, K. & Loukas, A. Implicitly guided design with propen: Match your data to follow the gradient. arXiv preprint arXiv:2405.18075 (2024).

24. Lin, Z. et al. Evolutionary-scale prediction of atomic-level protein structure with a language model. Science 379, 1123–1130 (2023).

25. Su, J. et al. Saprot: Protein language modeling with structure-aware vocabulary. bioRxiv 2023–10 (2023).

26. Hutchinson, M. et al. Toward enhancement of antibody thermostability and affinity by computational design in the absence of antigen. In mAbs, vol. 16, 2362775 (Taylor & Francis, 2024).

27. Starr, C. G. et al. Ultradilute measurements of self-association for the identification of antibodies with favorable high-concentration solution properties. Mol. Pharm. 18, 2744–2753 (2021).

28. Chai, Q., Shih, J., Weldon, C., Phan, S. & Jones, B. E. Development of a high-throughput solubility screening assay for use in antibody discovery. In MAbs, vol. 11, 747–756 (Taylor & Francis, 2019).

29. Honegger, A. & PluÈckthun, A. Yet another numbering scheme for immunoglobulin variable domains: an automatic modeling and analysis tool. J. molecular biology 309, 657–670 (2001).

30. Dunbar, J. & Deane, C. M. Anarci: antigen receptor numbering and receptor classification. Bioinformatics 32, 298–300 (2016).

31. Kovaltsuk, A. et al. Observed antibody space: a resource for data mining next-generation sequencing of antibody repertoires. The J. Immunol. 201, 2502–2509 (2018).

32. Suzek, B. E. et al. Uniref clusters: a comprehensive and scalable alternative for improving sequence similarity searches. Bioinformatics 31, 926–932 (2015).

33. Berman, H. M. et al. The protein data bank. Nucleic acids research 28, 235–242 (2000).

34. He, K., Zhang, X., Ren, S. & Sun, J. Deep residual learning for image recognition. In Proceedings of the IEEE conference on computer vision and pattern recognition, 770–778 (2016).

35. Paszke, A. et al. Pytorch: An imperative style, high-performance deep learning library. Adv. neural information processing systems 32 (2019).

36. Kingma, D. P. & Ba, J. Adam: A method for stochastic optimization. arXiv preprint arXiv:1412.6980 (2014).

37. Gainza, P., Nisonoff, H. M. & Donald, B. R. Algorithms for protein design. Curr. opinion structural biology 39, 16–26 (2016).

38. Luan, P. et al. Automated high throughput microscale antibody purification workflows for accelerating antibody discovery. In MAbs, vol. 10, 624–635 (Taylor & Francis, 2018).

39. Kabsch, W. xds. Acta Crystallogr. Sect. D: Biol. Crystallogr. 66, 125–132 (2010).

40. McCoy, A. J. et al. Ab initio solution of macromolecular crystal structures without direct methods. Proc. Natl. Acad. Sci. 114, 3637–3641 (2017).

41. Emsley, P. & Cowtan, K. Coot: model-building tools for molecular graphics. Acta crystallographica section D: biological crystallography 60, 2126–2132 (2004).

42. Adams, P. D. et al. Phenix: a comprehensive python-based system for macromolecular structure solution. Acta Crystallogr. Sect. D: Biol. Crystallogr. 66, 213–221 (2010).

43. Schrödinger, LLC. The PyMOL molecular graphics system, version 1.8 (2015). PyMOL The PyMOL Molecular Graphics System, Version 1.8, Schrödinger, LLC.

