## Supplemental File for "DyAb: sequence-based antibody design and property prediction in a low-data regime"

### Supplementary material

| | aEGFR lead | aEGFR-R1-1 |
|---|---|---|
| **PDB code** | 9MU1 | 9MSW |
| **X-ray source** | SSRF BL02U1 | SSRF BL02U1 |
| **Wavelength (Å)** | 0.97918 | 0.97918 |
| **Detector** | Eiger2 S 9M | Eiger2 S 9M |
| **Resolution range (Å)** | 47.61 – 2.43 | 202 - 2.10 |
| **Highest res. bin (Å)** | 2.53 – 2.43 | 2.14 - 2.10 |
| **Space group** | C2 | P32 |
| **Multiplicity** | 4.8, 4.5 | 6.9, 6.9 |
| **Completeness (%)** | 99.9, 100 | 100, 100 |
| **Mean $\mathbf{I}/\sigma_{\mathrm{I}}$** | 8.8, 2.2 | 9.9, 2.0 |
| **Wilson B ($\mathbf{Å^2}$)** | 34.7 | 25.9 |
| **CC $\frac{1}{2}$ (highest bin)** | 0.692 | 0.696 |
| **Rmerge (%)** | 19.5, 77.9 | 17.0, 113 |
| **# reflections ($\mathbf{R_{free}}$ set)** | 36,869 (1,909) | 159,121 (7,953) |
| **Resolution range (Å)** | 46.59 – 2.43 | 33.7 – 2.10 |
| **$\mathbf{R_{work}}$, $\mathbf{R_{free}}$ (%)** | 19.6, 24.8 | 17.8, 23.0 |
| **# non-H atoms** | 3,313 | 9,973 |
| **# solvent molecules** | 148 | 911 |
| **Rmsd bond lengths (Å)** | 0.004 | 0.004 |
| **Rmsd bond angles (°)** | 0.710 | 0.698 |
| **Ramachandran fav. (%)** | 96.3 | 98.0 |
| **Ramachand. outlier (%)** | 0.0 | 0.0 |
| **Ave B-factor ($\mathbf{Å^2}$)** | 41.0 | 31.4 |
| **Molprobity clash score** | 3.4 | 2.2 |

Table 1: Crystallographic data for aEGFR lead and top design

### Supplementary figures

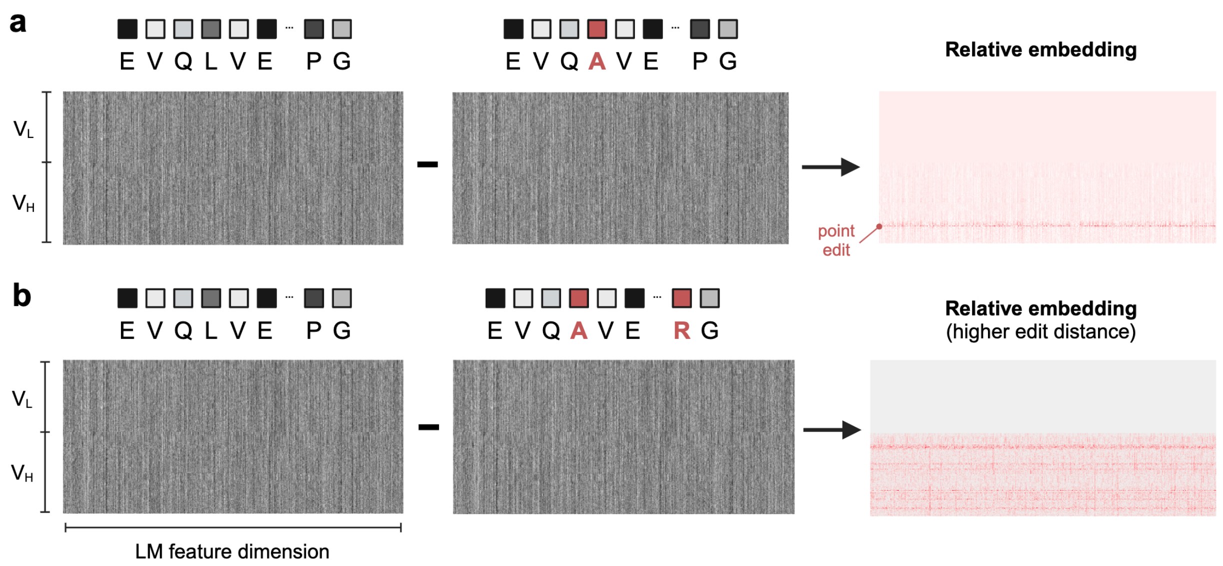


Figure S1: **Relative embeddings capture differences between pairs of protein sequences.** Example Antiberty embeddings for antibody variable domains that differ only by **(a)** a single residue and **(b)** eight residues in the heavy chain. The effects of these mutation(s) are captured by the relative embedding both near and far from the edit site (red, right).

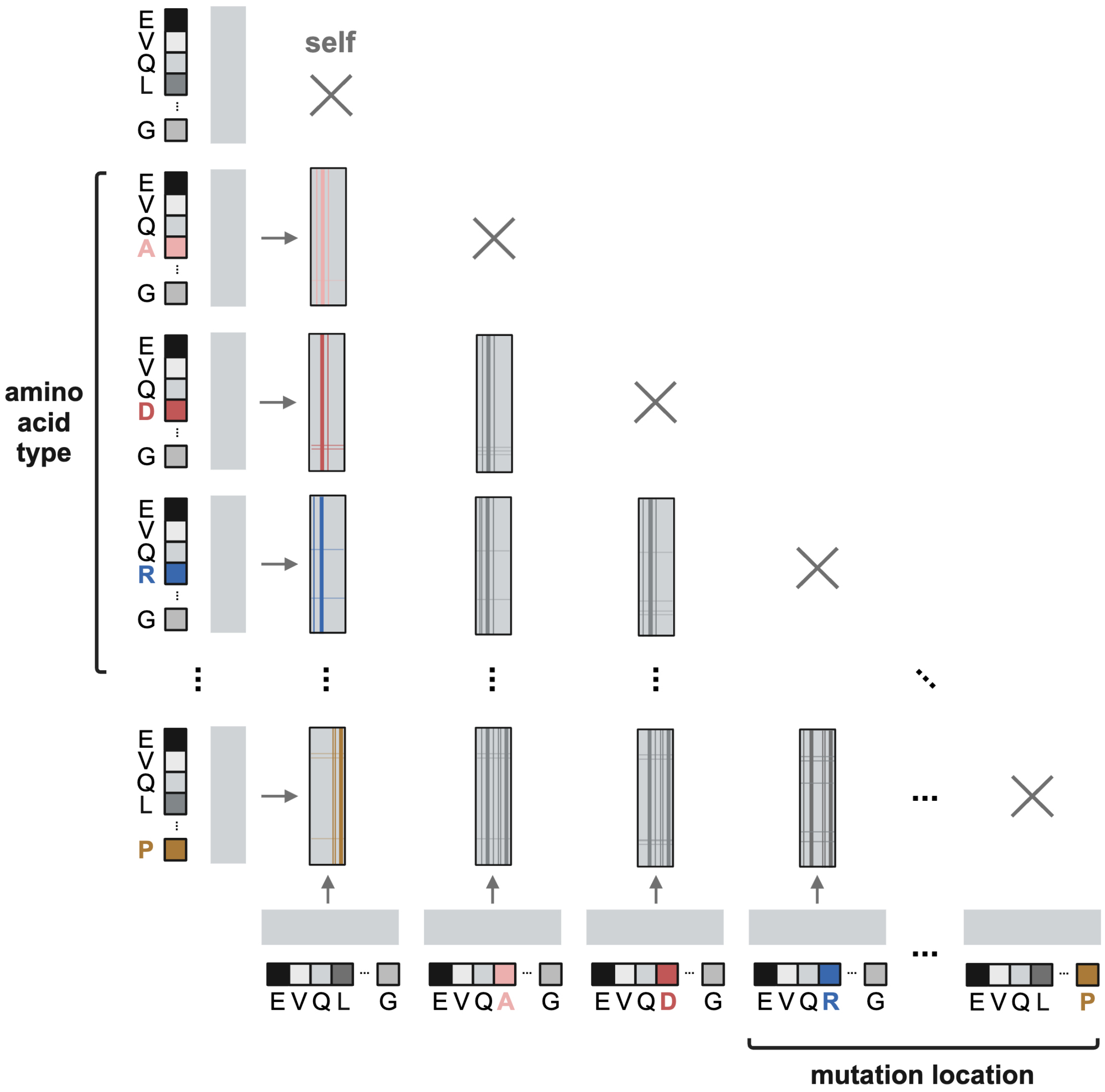


Figure S2: **Relative embeddings of point variants.** Cartoon schematic for a library of point variants that exhaustively sample mutation sites and amino acid types (labels). Relative embeddings capture differences between $\binom{n}{2}$ sequence pairs, rather than $n$ lone sequences.

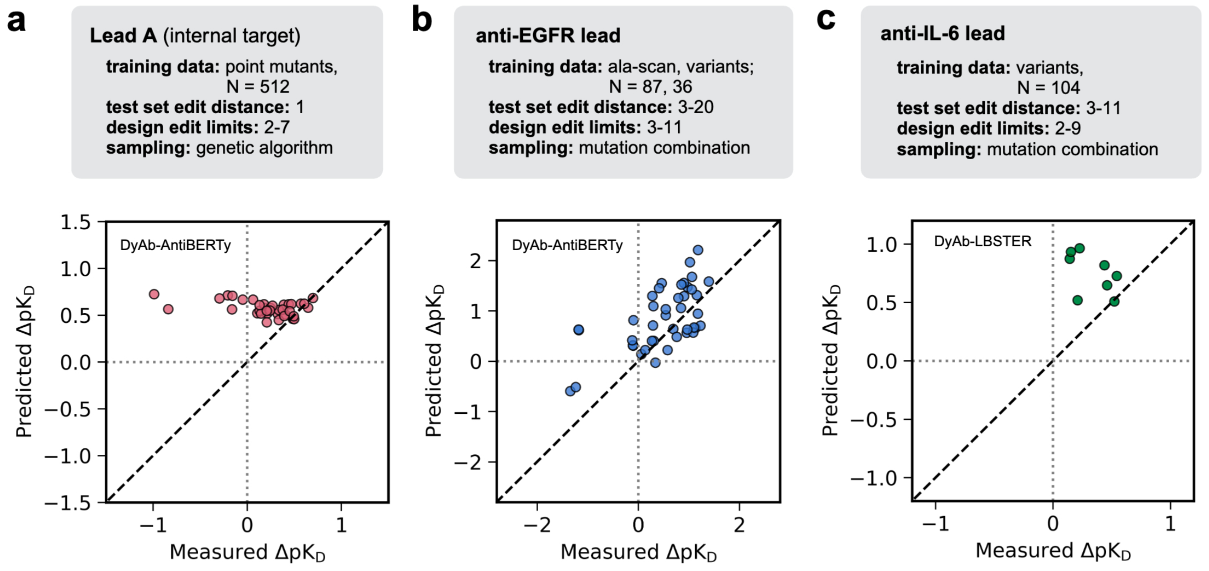


Figure S3: **DyAb model performance on tested designs.** Predicted versus measured improvements in affinity, $\Delta pK_D$, for variants of three lead antibodies (upper, colors, **(a-c)**), for the DyAb-generated designs selected for experimental testing. Affinity differences are computed relative to each lead molecule (grey dotted lines) by the trained DyAb models shown in Fig. 2.

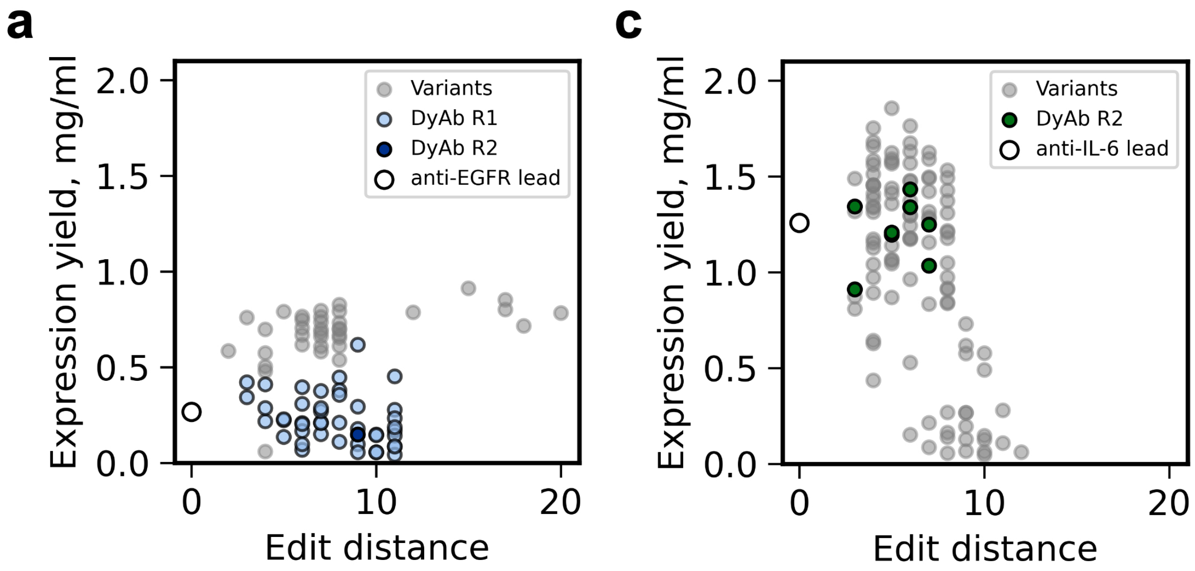


Figure S4: **DyAb designs maintain affinity and expression at higher edit distances.** Expression levels versus edit distance for variants in the training set (grey) and DyAb designs (colors) around the **(a)** anti-EGFR and **(b)** anti-IL-6 leads.

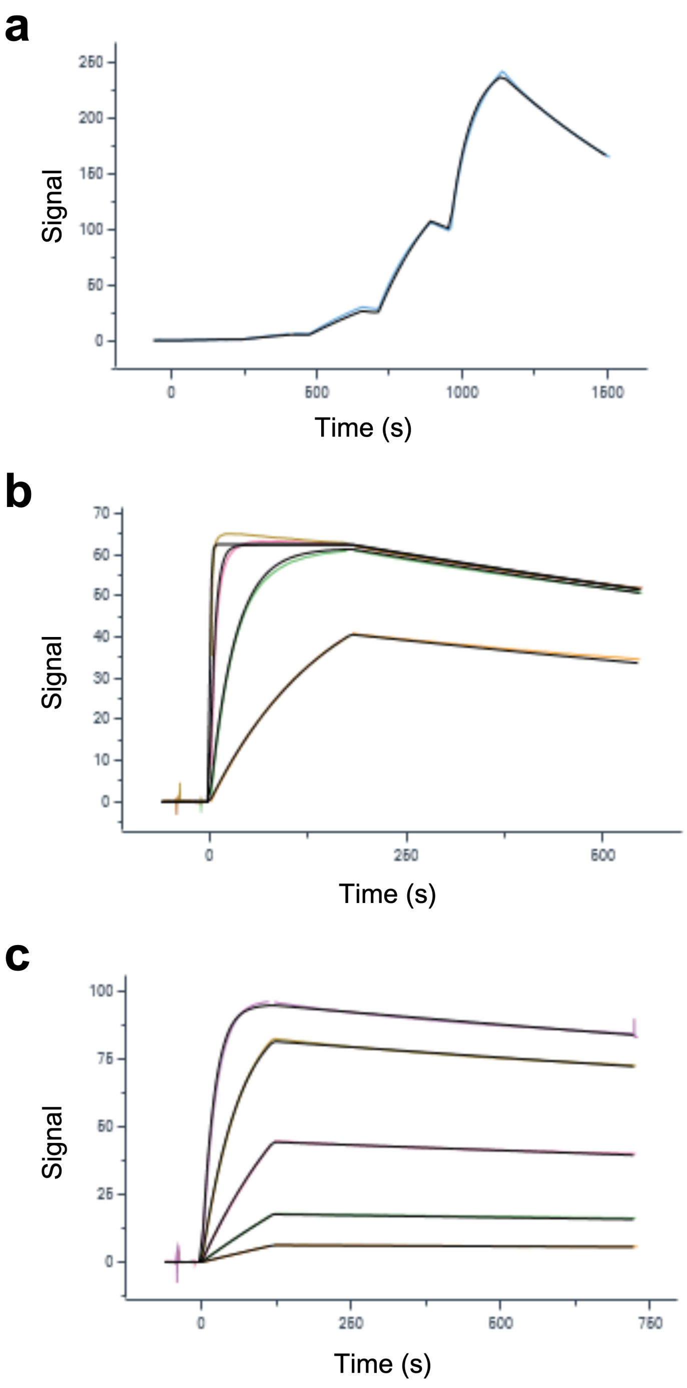


Figure S5: **Raw SPR curves for top DyAb designs.** Sensorgrams for the highest-affinity DyAb design for each seed, **(a)** seed A, **(b)** anti-EGFR and **(c)** anti-IL-6. The measurement mode was changed from single-cycle to multi-cycle kinetics in between testing for lead A and anti-EGFR.

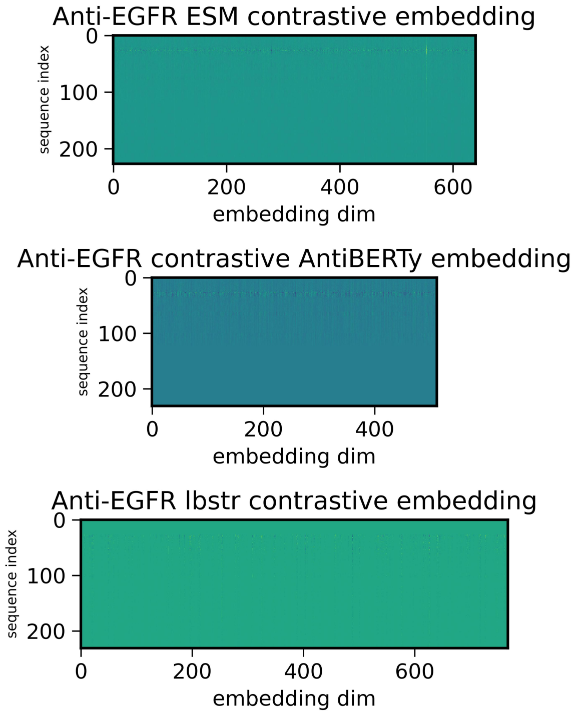


Figure S6: **Contrastive embedding of 2 anti-EGFR variants** Here we highlight the contrastive embedding of three different protein language model (AntiBERTy, ESM-2, LBSTER).

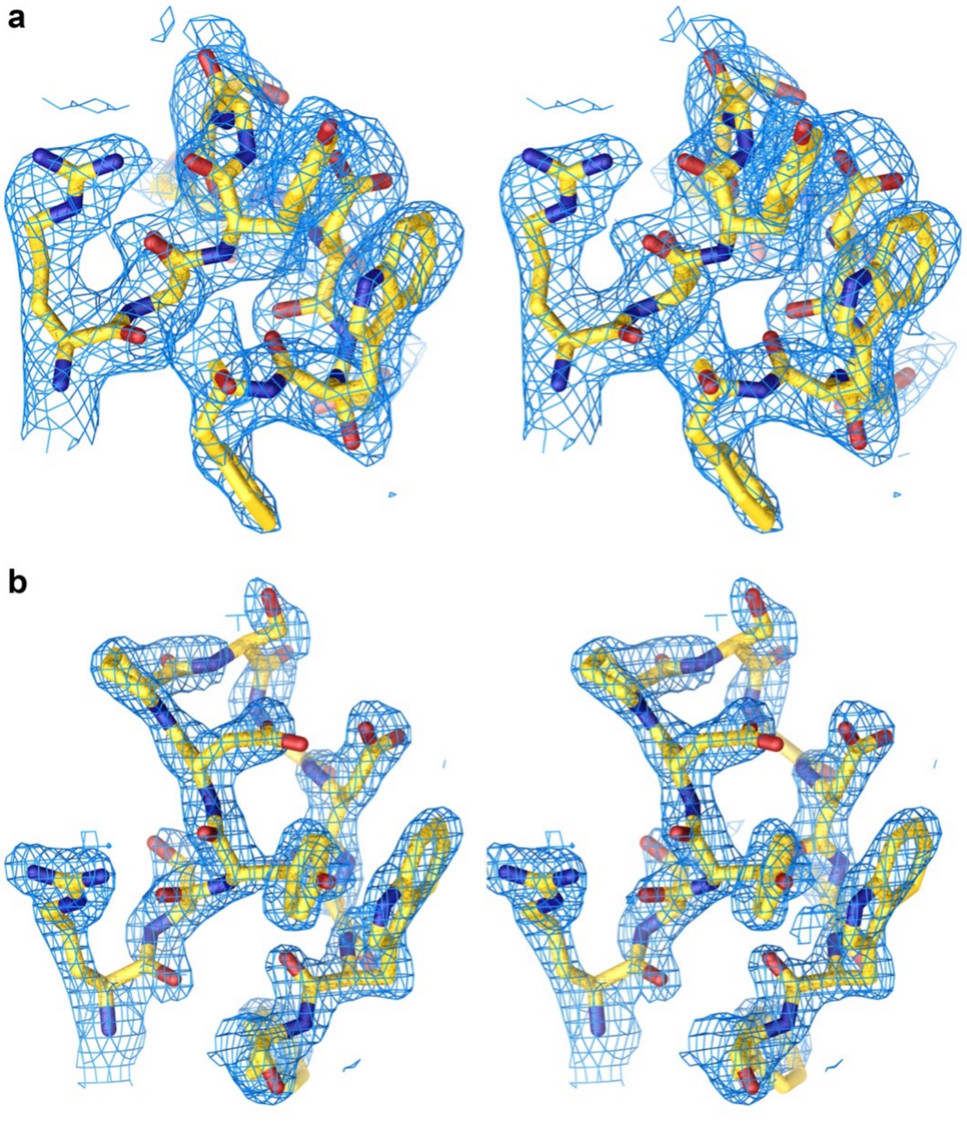


Figure S7: **Electron density maps.** Divergent eye stereo images of the simulated annealing composite omit difference electron density maps, $(2m|Fo| - D|Fc|)exp(i\alpha c)$, of Fabs for the aEGFR **(a)** lead, 9MU1, and **(b)** R1-1 design, 9MSW, contoured at $1\sigma$. The region shown is the heavy chain CDR-H3 loop, from residue Arg94-Phe100D (Kabat numbering).
